# Virtual reality simulation of epi-retinal prosthetic vision highlights the relevance of the visual angle

**DOI:** 10.1101/2020.07.18.195800

**Authors:** Jacob Thomas Thorn, Enrico Migliorini, Diego Ghezzi

**Author notes:** Correspondence and requests for materials should be addressed to D.G.

## Abstract

**Objective:** Retinal prostheses hold the potential to restore artificial vision in blind patients suffering from outer retinal dystrophies. The optimal number, density, and coverage of the electrodes that a retinal prosthesis should have to provide adequate artificial vision in daily activities is still an open question and an important design parameter needed to develop better implants.

**Approach:** To address this question, we investigated the interaction between the visual angle, the pixel number and the pixel density without being limited by a small electrode count, as in previous research reports. We implemented prosthetic vision in a virtual reality environment in order to simulate the real-life experience of using a retinal prosthesis. We designed four different tasks simulating: object recognition, word reading, perception of a descending step and crossing a street.

**Main results:** The results of our study showed that in all the tasks the visual angle played the most significant role in improving the performance of the participant.

**Significance:** The design of new retinal prostheses should take into account the relevance of the restored visual angle to provide a helpful and valuable visual aid to profoundly or totally blind patients.

## Introduction

Retinal prostheses are neuroprostheses able to revert blindness caused by outer retinal dystrophies, such as retinitis pigmentosa and dry age-related macular degeneration[1,2]. Over two decades, retinal prostheses yielded extraordinary results in patients[3–5]; however, despite the progresses made in the research field, there are still several limitations affecting the possibility to provide useful vision in daily life[6]. Blindness is defined as visual acuity of less than 20/400 or a corresponding visual field loss to less than 10 degrees, in the better eye with the best possible correction (World Health Organization). In North America and most of European countries, legal blindness is defined as visual acuity of 20/200 or visual field no greater that 20 degrees.

Although visual acuity is an important measure of the quality of vision (both natural and artificial), another very important feature is the size of the visual field, which might affect several visually-guided behaviours important in daily life. For retinal prostheses, as a rule of thumb, the visual field is linked to the retinal coverage of the prosthesis, while the visual acuity to the density of the stimulating electrodes. Therefore, the number, density, and coverage of the electrodes is an important design parameter for retinal prostheses to provide adequate artificial vision in daily life. The retinal prostheses in clinical use are limited by the number of pixels that can be wired to the implantable stimulator, which impose a small retinal coverage and/or a low pixel density. As a consequence, the clinical devices were intrinsically limited to either increase the pixel density in a small surface area[3] or enlarge the surface area at the expense of the pixel density[4,7].

An early study on healthy participants under pixelated vision indicated that an array of 25 × 25 pixels and 30 degrees of visual angle could provide adequate mobility skills[8]. It is worth noting that, combined together, these parameters (625 pixels and 30 degrees) are beyond what is offered by retinal prostheses in clinical use today. A follow up study confirmed the results reporting that a visual angle of 33 × 23 degrees was preferred by participants for mobility with more that 1,000 pixels to allow for safer decision-making[9]. Other studies estimated that, to be useful in daily life, a retinal prosthesis should have at least 500 pixels in the central area of about 5 mm in diameter[10,11]. More recently, a trial on healthy participants showed that the number of pixels required to recognize common objects is on the order of 3,000 to 5,000[12]. Collectively, these results suggest that covering a large visual angle even with a limited resolution seems sufficient for mobility skills, while high pixel density is required for object recognition. Unfortunately, all of these previous studies focused on only one part of the problem, either the wide visual angle or the high pixel density. In addition, these studies were often based on a conversion of images into an idealized phosphene representation, where phosphenes are round and perfectly aligned. Although this might hold true for sub-retinal prostheses, this is not the case for epi-retinal devices. Due to the direct activation of axons in the nerve fibre layer, phosphenes are often elongated or with a complex shape[13,14]. Since, wide-field arrays have been so far designed for epiretinal placement only, the complex shape of phosphenes must be considered when simulating artificial vision.

Recently, photovoltaic retinal prostheses allowed for wireless retinal stimulation[15–22], thus overcoming in principle the problem of prioritizing between pixel density or surface area. Photovoltaic retinal prostheses have the ability to embed a large number of pixels, limited only by the manufacturing capability and the conversion efficiency of the pixel. Moreover, conformable wide-field retinal prostheses enabled the enlargement of the retinal coverage without affecting the pixel density[23].

These research results prompted us to investigate the interaction between the retinal coverage and the pixel density without being limited to a small number of electrodes. We took advantage of virtual reality (VR) software paired with a portable head mounted display (HMD) to monitor the performance of healthy participants under simulated prosthetic vision with variable pixel density and retinal coverage. The stimulation layout was mapped accordingly to the epi-retinal wide-field retinal prosthesis known as POLYRETINA[23], which allows in the same device both a high pixels density and a wide visual field. Since POLYRETINA is an epi-retinal prosthesis, the direct activation of axons resulting in elongated phosphenes was considered according to recent computational models[13]. Also, taking advantage of the flexibility allowed by VR, we tested behavioural performances not only for word and object recognition, but also in common daily activities, such as recognising a descending pair of steps or crossing a busy street.

Our results provide insights about the optimal number, density, and coverage required for a retinal prosthesis to be useful in daily life. In particular we found that the visual angle had the most significant effect in improving the performance of the tested participant. These results led to the conclusion that in order to be useful in daily life, retinal prostheses must restore artificial vision in a large visual angle so as to overcome limitations related to head movements for space scanning and mental reconstruction of the visual scene.

## Methods

### Experiment design

The participants performed four behavioural tests, described as ‘object recognition’, ‘word reading’, ‘step perception’, and ‘street crossing’.

The tests were performed using a Dell Precision 3630 desktop computer with an Intel Xeon E-2146G CPU (3.50 GHz) and an nVidia GeForce GTX 1080 GPU. To simulate prosthetic vision two commercial VR HMDs were used: the FOVE (‘object recognition’ and ‘word reading’) and the VIVE Pro Eye (‘step perception’ and ‘street crossing’). The FOVE has a resolution of 2560 × 1440 pixels (1280 × 1440 pixels per eye), covers a visual field of approximately 95 degrees, and updates at 70 Hz. The HMD collects data about the participant’s head orientation through internal gyroscopes, and tracks eye gaze through two integrated infrared eye-tracking cameras at a frequency of 120 Hz. Correspondingly, the VIVE Pro Eye has a resolution of 2880 × 1600 (1440 × 1600 pixels per eye), a maximum field of view of 110 degrees and a 90Hz refresh rate. Head orientation and position is provided using two external cameras aimed at the headset, and eye-tracking is similarly accomplished using integrated infrared cameras operating at 120 Hz.

During the ‘object recognition’ and ‘word reading’ tests, the participants were seated, wearing the FOVE HMD, and holding a game controller. Participants were then shown a series of objects or words and asked to identify them. When the participants felt like they had recognized the object or word, they would press a button on the game controller and then tell their guess to the examiner, who would mark the answer as correct or incorrect and proceed to the next item (minimum time interval of 1s). Participants were allowed to explore the objects or words via head movements and eye movements. For the objects, participants also had the ability to rotate the object with the game controller. Words were either presented in English, French or Italian at the preference of the participant from a list of commonly used words. The full list of 108 possible objects is shown in **Fig. 1**.

**Figure 1.**
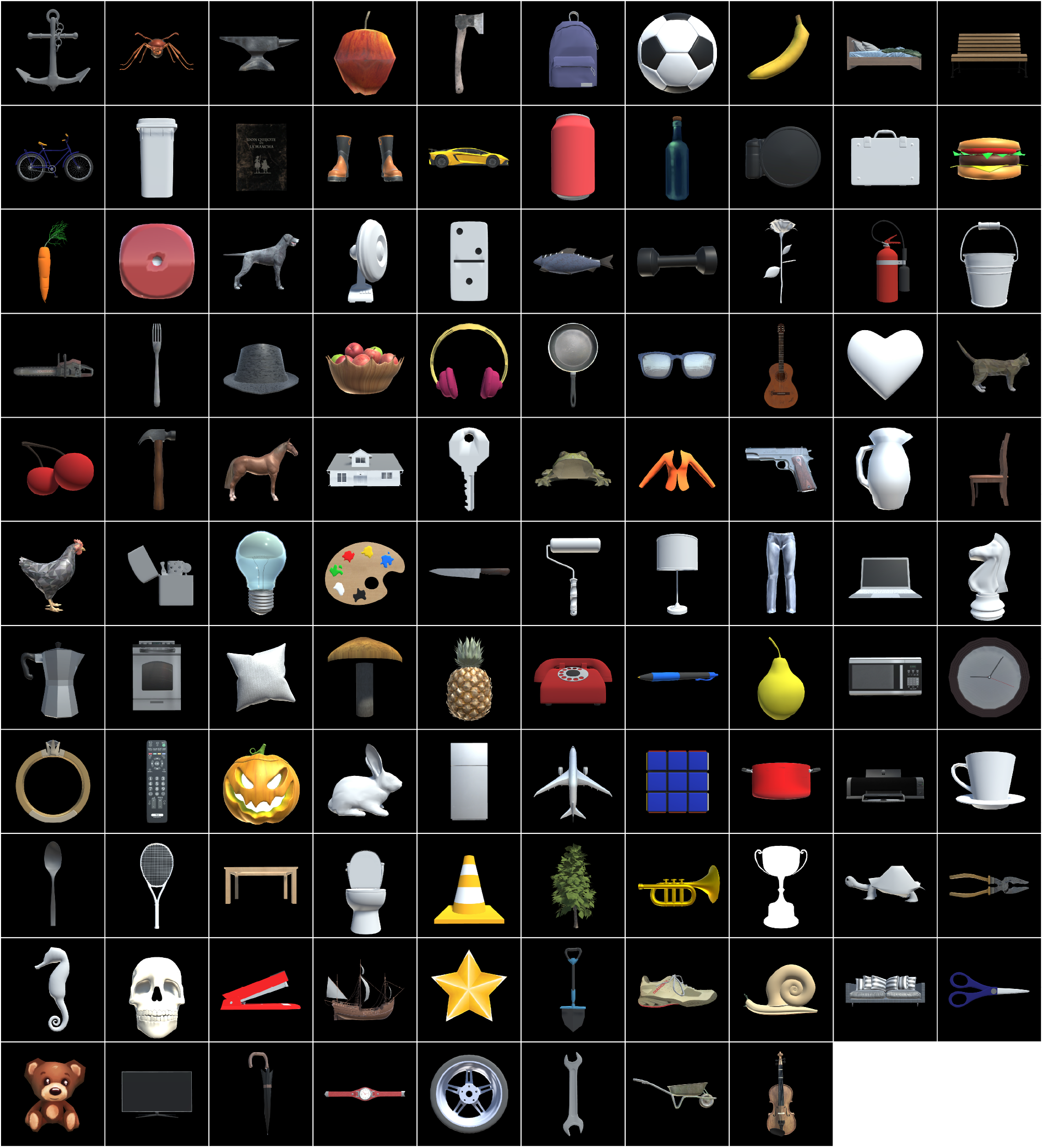
Images of the 108 common objects used during the ‘object recognition’ test.

During the ‘step perception’ test, participants were asked to stand while wearing the VIVE Pro Eye HMD and holding two VR controllers. The virtual scenario involved the participants standing on a platform with two steps placed diagonally in front of them. The height of both steps was altered in each trial and participants were instructed to determine which step they thought was highest, or the easiest to step down on to. Participants were allowed to explore the environment via head and eye movements. Once they had made a decision, they would press a button on the corresponding left or right controller. Differences in height were 10, 20, 30 and 40 cm.

During the ‘street crossing’ test, the virtual environment replicated a trafficked street, with cars moving at a speed of 30 km hr^−1^ in both directions, and the participants standing on the edge of the sidewalk. Participants were allowed to explore the environment via head and eye movements. At random intervals between 8 and 12 seconds, there would be a gap of 2, 4, 6, or 8 seconds on the left side of the street and 4, 6, or 8 seconds on the right (note that the length of the opening on the two sides could be different) where no cars would be present. If the participants felt that it was safe to cross the street, they would take two steps forward. The program would then determine that the participant had crossed the road, based on data received from the VIVE motion capture system. In case there was a car too close to the participant’s position, the system would mark the crossing as failed and play the sound of a car crashing, otherwise the crossing would be marked as successful. In either case, the participant would then return to the starting position and a new layout would be loaded. The length of the openings determined if the crossing would be impossible (*), possible but requiring a quick decision (**), or relatively easy (***), as in **Tab. 1**.

**Table 1.** Gaps combination in the ‘street crossing’ test. Values are in seconds.

|  |  |  |  |  |
| --- | --- | --- | --- | --- |
|  |  | Gap on the right |  |  |
|  |  | 4 | 6 | 8 |
| Gap on the left | 2 | * | * | * |
|  | 4 | ** | *** | *** |
|  | 6 | ** | *** | *** |
|  | 8 | ** | *** | *** |

### Simulation of prosthetic vision

All of the test environments as well as the simulation of prosthetic vision were developed using Unity (www.unity.com). Data collection was also performed in Unity. For tests using the FOVE headset, eye-tracking information was obtained via the FOVE Software Development Kit (SDK). Otherwise, when using the VIVE Pro Eye HMD, the SRanipal eye-tracking SDK was used to track participants eye.

The simulated prosthetic vision was computed using Cg shaders, allowing the system to run in real-time. Briefly, shaders are pieces of code that are run on a computer’s graphics hardware for each output pixel (FOVE:1280×1440; VIVE Pro Eye: 1440×1600), every update frame (FOVE: 75Hz; VIVE Pro Eye: 90Hz). To save on computing time, phosphene centres were pre-calculated and saved as a texture which could be accessed by the shaders. In this sense, the texture equated to a data matrix with a width and height equal to the resolution of the output image, where each element in the matrix held the coordinate of the centre of the closest phosphene to the pixel. Pixels were then activated in the Cg shader depending on the following criteria: a) they are inside the POLYRETINA field of view, b) they are inside their closest phosphene, c) their closest electrode is not “broken” (10% chance), and d) the luminance of the input image at the equivalent pixel exceeded 0.5. Images were converted into phosphenes using four layouts based on the POLYRETINA prosthesis: 100/150, 80/120, 60/90, and 40/60 (electrode diameter in μm / electrode pitch in μm). The number of pixels for each layout is reported in **Tab. 2** as a function of the visual angle.

**Table 2.** Number of phosphenes for each layout.

| Table 2. Number of phosphenes for each layout. |  |  |  |  |
| --- | --- | --- | --- | --- |
| Visual angle | Layout 1: 100/150 | Layout 2: 80/120 | Layout 3: 60/90 | Layout 4: 40/60 |
| 5° | 86 | 119 | 214 | 475 |
| 10° | 355 | 477 | 855 | 1,932 |
| 15° | 800 | 1,088 | 1,942 | 4,374 |
| 25° | 2,244 | 3,045 | 5,437 | 12,241 |
| 35° | 4,417 | 5,995 | 10,688 | 24,073 |
| 45° | 7,321 | 9,914 | 17,681 | 39,814 |

For each phosphene image, three sources of random variability were included: the phosphene size was randomly varied between −30% to +30% of the electrode size, the phosphene brightness was randomly varied between 50% (grey) to 100% (white) of the phosphene’s default brightness, and 10% of the electrode were considered not functional. Lastly, image presentation was not continuous but pulsed. Ideally, we would have planned 10-ms image presentation followed by 40-ms of dark (equivalent to a 20 Hz repletion rate). However, due to the refresh rate of both the FOVE and VIVE Pro Eye being higher than 10 ms (13 ms for the FOVE and 11 ms for the VIVE Pro Eye), the black pause was 39 ms for the FOVE and 44 ms for the VIVE Pro Eye.

The process just described generates a phosphene image, often called a scoreboard (SB) model. However, based on the report from participants implanted with epi-retinal prostheses, phosphenes are more likely to be interpreted as curved oblong-like shapes[13]. A second Cg shader was used to account for the axonal fibre activation, having as an input the phosphene image. The values describing the trajectory of the axonal fibre passing through each pixel were pre-generated using a mathematical model[24] to reduce the amount of real-time processing. The shader then used these values to generate each phosphene’s tail, following the underlying axon fibres. The tail’s brightness would dissipate as a function of the distance from its phosphene, using the activation function described in [25]. Finally, the tails were superimposed on top of the original phosphene image. This image representation is called an axon map (AM) model. After this, one final Cg shader applies a small 3-pixel Gaussian blur to the image in order to soften the pixel edges for both the SB and SM models. The image is then displayed in the right eye only of the VR headset.

For the ‘object recognition’, ‘step perception’ and ‘street crossing’ tests, an edge detection algorithm was also applied (for both the SB and AM model) just before the phosphene shader. The implementation is a modified version of the widely used Canny edge detection algorithm[26], except that instead of thinning the edges, it was preferable to thicken them to ensure that enough phosphenes were activated to represent the detected edge. The entire code of the project is available online at https://github.com/lne-lab/polyretina_vr.

### Ethical statement

Experiments were approved by the human research ethics committee of École polytechnique fédérale de Lausanne (decision number 042-2018 / 16.10.2018). Seventeen healthy volunteers were involved in the study (**Tab. 3**). However, due to logistical reasons not all the participants performed all the tests.

| Participant | Sex | Age | Mother tongue | Fig 2 | Fig 3 | Fig 4 | Fig 6 | Fig 7 | Fig 8 |
| --- | --- | --- | --- | --- | --- | --- | --- | --- | --- |
| 1 | Female | 25 | Italian | x |  | x |  |  |  |
| 2 | Female | 29 | French | x |  | x |  |  |  |
| 3 | Male | 26 | Italian | x |  | x |  |  |  |
| 4 | Female | 28 | French | x |  | x |  |  |  |
| 5 | Male | 38 | Italian |  | x |  | x | x |  |
| 6 | Female | 31 | French |  | x |  | x | x |  |
| 7 | Female | 25 | French |  | x |  | x | x | x |
| 8 | Female | 28 | Italian |  | x |  | x | x | x |
| 9 | Female | 29 | Italian |  | x |  | x | x | x |
| 10 | Male | 28 | French |  | x |  | x | x | x |
| 11 | Female | 25 | Italian |  | x |  | x | x | x |
| 12 | Male | 28 | French |  | x |  | x |  |  |
| 13 | Female | 25 | Italian |  | x |  | x | x | x |
| 14 | Female | 22 | French |  | x |  | x |  |  |
| 15 | Female | 25 | English |  |  |  |  | x | x |
| 16 | Male | 24 | Italian |  |  |  |  | x | x |
| 17 | Female | 33 | Farsi |  |  |  |  | x | x |

## Results

### Object recognition

The first test assesses the capability to recognize common objects (35 ° in size) under prosthetic vision (**Fig. 1**).

In a first cohort of participants (*n* = 4; participants 1 to 4), images were converted using the SB model and the layout with the lowest resolution (100/150). The participant’s visual field was restricted to various visual angles: 5, 10, 15, 25, 35, and 45 ° (**Fig. 2a**). For each visual angle, 18 objects were randomly presented (108 objects per participant). We evaluated the success rate as percentage of correct answers (**Fig. 2b**) and the time required to provide a correct answer (**Fig. 2c**). In the latter, the time was not considered when the given answer was not correct. Both measures showed that the increase of the visual angle resulted in a higher success rate and lower time to response. For each participant, we quantified a performance index to account for both the success rate and the time taken to provide a correct answer[27,28], defined as in equation (1):

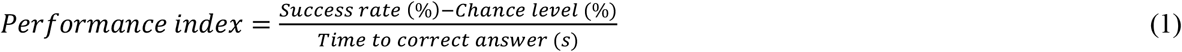

**Figure 2.**
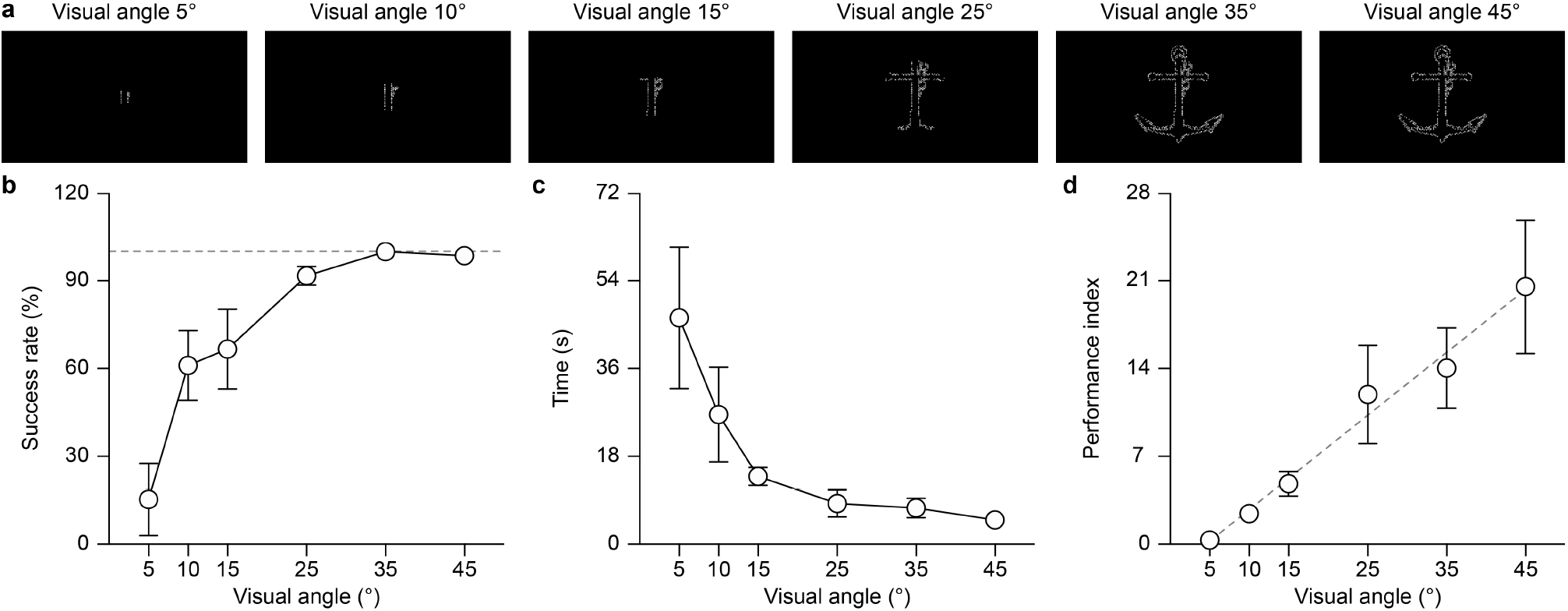
(a) Rendering of one object with the SB model and the 100/150 layout as a function of the visual angle. (b) Success rate as a function of the visual angle (mean ± s.d., *n* = 4 participants). The grey dashed line shows a success rate of 100%. (c) Time to provide a correct answer as a function of the visual angle (mean ± s.d., *n* = 4 participants). (d) Performance index as a function of the visual angle (mean ± s.d., *n* = 4 participants). The grey dashed line shows the linear regression model.

For the ‘object recognition’ experiment, the chance level was 0 since the objects were unknown. A linear regression model showed that the performance index (**Fig. 2d**) increased with the visual angle (slope = 0.50; R squared = 0.87) and the slope is significantly non-zero (p < 0.0001, F = 141.1, DFn = 1, DFd = 22). This result highlights the important role of the visual angle in retinal prostheses.

The SB model is a simplified representation of the visual perceptions occurring with epi-retinal prostheses. Therefore, in a second cohort of participants (*n* = 10, participants 5 to 14), we implemented the AM model which accounts for the activation of the fibres of passage. We tested the impact of the pixel layout and the tail length on the capability to recognise objects. The visual field of the participant was kept unconstrained (45 °), to highlight only the effect of the above-mentioned parameters. Four layouts (100/150, 80/120, 60/90, and 40/60) and three tail lengths (1, 2, 3 °) were randomly varied to obtain a total of 12 possible configurations (**Fig. 3a**). For each configuration 9 different objects were randomly presented (108 different objects per participant). As before, we evaluated the success rate (**Fig. 3b**) and the time required to provide a correct answer (**Fig. 3c**). From those parameters we estimated the performance index (**Fig. 3d**). For each tail length, the linear regression model revealed that the slopes are not significantly different than zero, thus suggesting that the layout had not a relevant role once the visual angle was maximised (**Tab. 4**).

**Table 4.** Result of the linear regression model: y = B_0_ + B_1_x.

| Table 4. Result of the linear regression model: $y = B_0 + B_1x$. | | | |
| --- | --- | --- | --- |
|  | Tail length: 1° | Tail length: 2° | Tail length: 3° |
| Slope / $B_1$ | 2.293 | 0.8139 | 1.182 |
| R squared | 0.04905 | 0.01034 | 0.02214 |
| p value | 0.1696 | 0.5324 | 0.3595 |
| F | 1.9600 | 0.3971 | 0.8604 |
| DFn | 1 | 1 | 1 |
| DFd | 38 | 38 | 38 |

**Figure 3.**
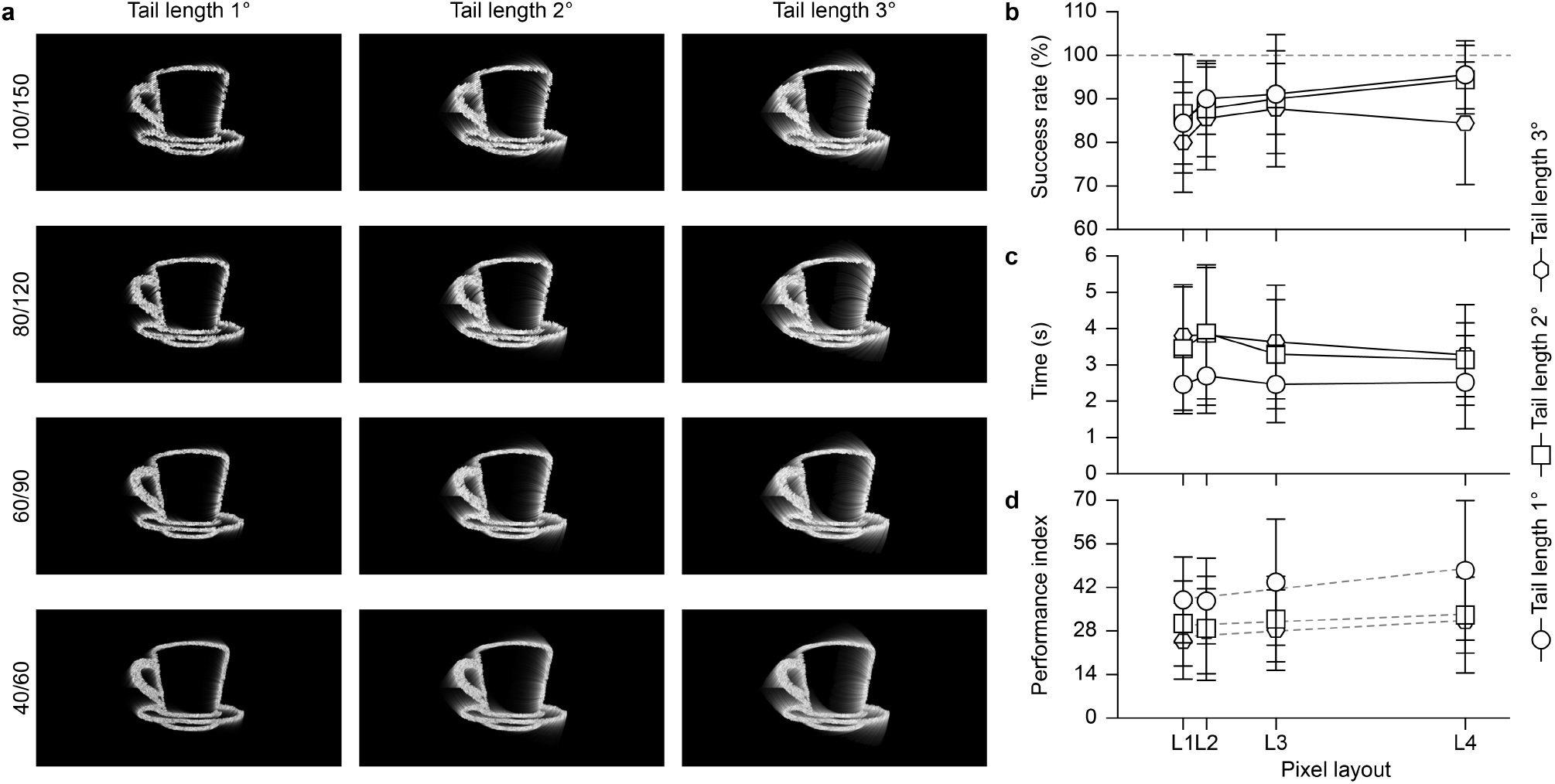
(a) Rendering of one object with the AM model as a function of the tail length (columns) and the pixel layout (rows). (b) Success rate as a function of the pixel layout for the three tail lengths (mean ± s.d., *n* = 10 participants). The grey dashed line shows a success of rate 100%. (c) Time to provide a correct answer as a function of the pixel layout for the three tail lengths (mean ± s.d., *n* = 10 participants). (d) Performance index as a function of the pixel layout for the three tail lengths (mean ± s.d., *n* = 10 participants). The grey dashed lines show the linear regression model. In panels (b-d) the x-axis was scaled according to the proportion of the number of pixels for each layout: 1 for L1 (layout 1, 100/150), 1.36 for L2 (layout 2, 80/120), 2.42 for L3 (layout 3, 60/90), and 5.32 for L4 (layout 4, 40/60).

Polling the data together, the differences between the slopes (obtained with the three tail lengths) are also not statistically significant (p = 0.7430, F = 0. 2978, DFn = 2, DFd = 114). Since the slopes are not significantly different, it is possible to calculate one slope for all the data (B_1_ = 1.430). However, the line elevations (B_0_) showed a statistically significant difference (p = 0.0002, F = 9.105, DFn = 2, DFd = 116) suggesting that the tail length affected the performance index.

The ‘object recognition’ experiment revealed that the pixel layout had little or no effect. On the other hand, the performance is highly dependent on the visual angle. The tail length also affected object recognition, with longer tails reducing the performance.

### Word reading

An advantage of a large visual field is the possibility to display entire words in the patient’s field of view instead of single letters. This second test assesses the capability to read words under wide-field prosthetic vision. A list of words composed by 4 to 6 letters was generated in the mother language of each participant and presented with the AM model (tail lengths: 1, 2, 3 °).

In the first cohort of participants (*n* = 4, participants 1 to 4), images were converted using the 100/150 layout and the letter height was fixed to 7 °, thus each word occupied a visual field ranging from 15.6 ° (4 letters) to 40.6 ° (6 letters). The participant’s visual field was restricted to various visual angles: 5, 10, 15, 25, 35, and 45 °. For each configuration (visual angle and tail length) 10 different words were randomly presented for a total of 180 different words per participant (**Fig. 4a**). Similar to the ‘object recognition’ test, both the success rate (**Fig. 4b**) and the time required to provide a correct answer (**Fig. 4c**) showed that the increase of the visual angle resulted in a higher success rate and lower time to response. As a consequence, the performance index also increased with the visual angle (**Fig. 4d**). For the ‘word reading’ experiment, the chance level was 0 since the words were unknown. For each tail length, a nonlinear regression model (second order polynomial) revealed that the second order polynomials are significantly non-zero (B_1_ and B2 coefficients) thus suggesting that the visual angle significantly affected the performance index (**Tab. 5**). Polling the data together, one curve cannot fit all the data sets (p < 0.0001, F = 5.893, DFn = 6, DFd = 63) suggesting that the tail length also affected the performance index.

**Table 5.** Result of the nonlinear regression model: y = B_0_ + B_1_x + B_2_x^2^.

| Table 5. Result of the nonlinear regression model: $y = B_0 + B_1x + B_2x^2$. | | | | | | | | | |
| --- | --- | --- | --- | --- | --- | --- | --- | --- | --- |
|  | Tail length: 1° |  |  | Tail length: 2° |  |  | Tail length: 3° |  |  |
| R squared | 0.7340 |  |  | 0.8752 |  |  | 0.7823 |  |  |
| | $B_0$ | $B_1$ | $B_2$ | $B_0$ | $B_1$ | $B_2$ | $B_0$ | $B_1$ | $B_2$ |
| Best-fit values | -6.022 | 2.789 | -0.03632 | -5.929 | 2.385 | -0.03213 | -4.671 | 2.116 | -0.3119 |
| p value | 0.3625 | 0.0003 | 0.0099 | 0.0865 | < 0.0001 | < 0.0001 | 0.1992 | < 0.0001 | 0.0002 |
| F | 0.8655 | 18.32 | 8.051 | 3.235 | 51.56 | 24.25 | 1.757 | 35.54 | 20.02 |
| DFn | 1 | 1 | 1 | 1 | 1 | 1 | 1 | 1 | 1 |
| DFd | 21 | 21 | 21 | 21 | 21 | 21 | 21 | 21 | 21 |

**Figure 4.**
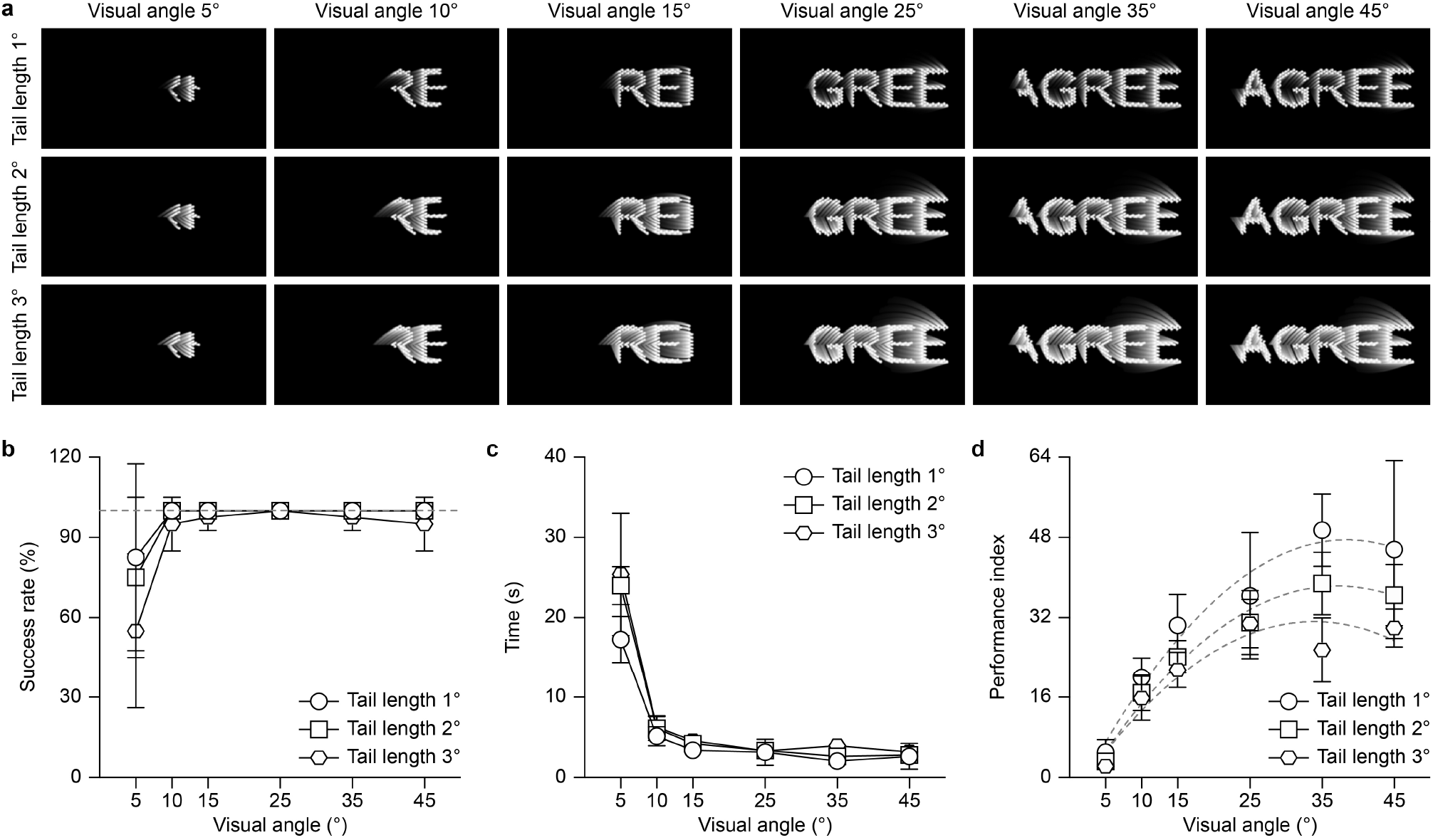
(a) Rendering of one word (letter heights 7 °) with the axon map model as a function of the visual angle (columns) and the tail length (rows). (b) Success rate (mean ± s.d., *n* = 4 participants) as a function of the visual angle for the three tail lengths. The grey dashed line shows a success rate of 100%. (c) Time to provide a correct answer (mean ± s.d., *n* = 4 participants) as a function of the visual angle for the three tail lengths. (d) Performance index (mean ± s.d., *n* = 4 participants) as a function of the visual angle for the three tail lengths. The grey dashed lines show the nonlinear regression model (second order polynomial).

The visual angle is a key parameter to improve the performance in reading words, so in the second cohort of participants (*n* = 10, participants 5 to 14), we tested the combined impact of the pixel layout, the tail length, and the letter height on the word recognition capability. The visual angle of the participant was kept unconstrained (45 °), to highlight only the effect of the above-mentioned parameters. The underlying hypothesis is that with a wide visual angle, the same word can be enlarged to maximise its readability. Five letter heights (from 3 ° to 7 °), four layouts (100/150, 80/120, 60/90, and 40/60), and three tail lengths (1, 2, and 3 °) were randomly varied to obtain a total of 60 possible configurations (**Fig. 5**). The words occupied an average visual field ranging from 6.4 ° (4 letters, 3 ° height) to 40.6 ° (6 letters, 7 ° height). For each configuration 10 different words were randomly presented for a total of 600 different words per participant. The experiment was split in three session of 200 words each to limit fatigue.

**Figure 5.**
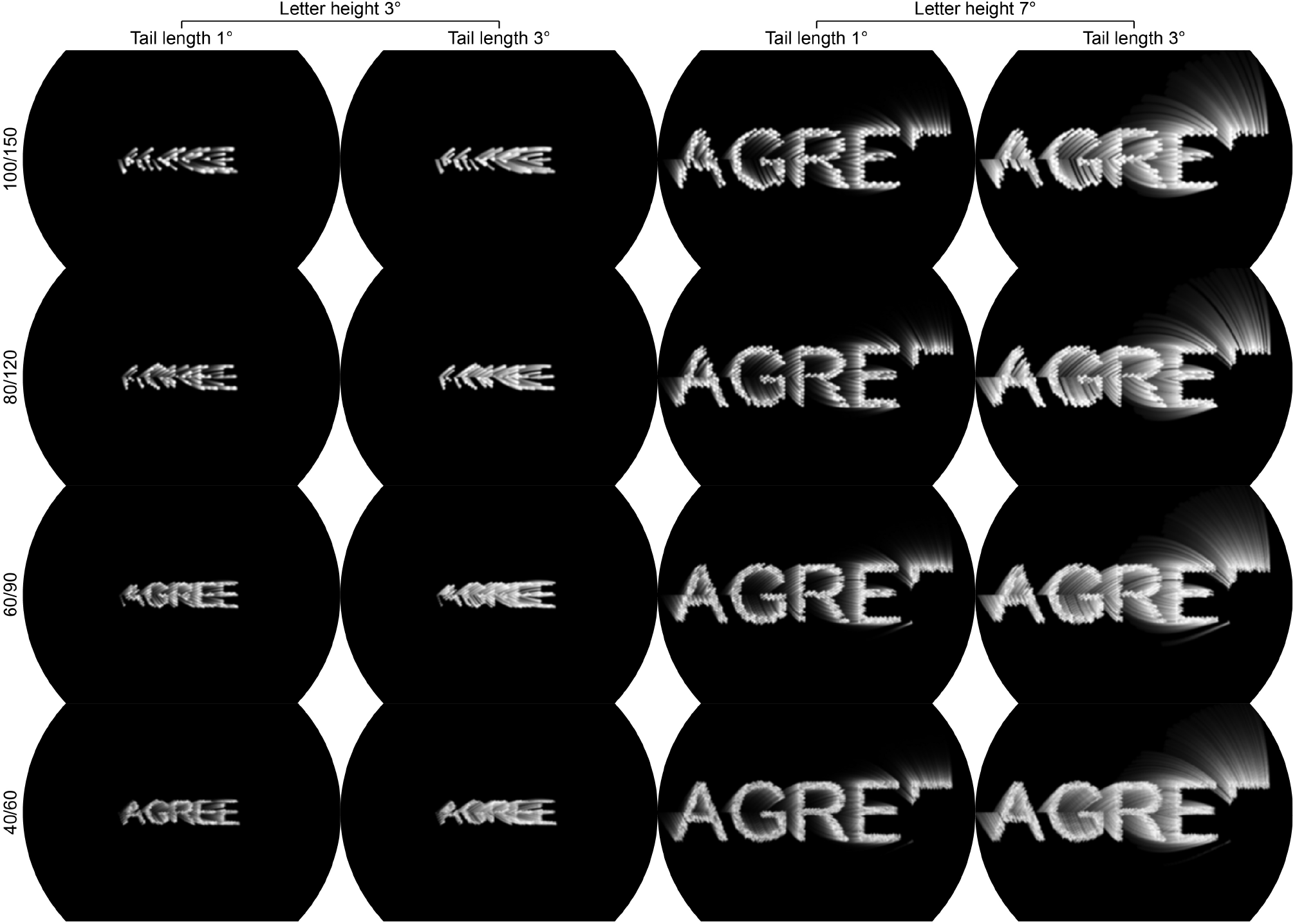
Rendering of one word with the AM model as a function of the tail length (columns) and the pixel layout (rows). Only the two extreme letter heights (3 ° and 7 °) and the two extreme tail length (1 ° and 3 °) are shown.

Qualitatively, for all the three tail lengths considered, both the success rate (**Fig. 6a**) and the time required to provide a correct answer (**Fig. 6b**) improved by increasing the pixel density and the letter height, in particular for the smallest letter heights (3 ° and 4 °) and the sparser layouts (100/150 and 80/120). To quantitively estimate the effect of the pixel layout, we split the dataset in three groups based on the tail length and performed linear and nonlinear regression on the performance index (**Fig. 6c-e**). For each tail length, the two sparser layouts (100/150 and 80/120) were fitted by a nonlinear regression model (second order polynomial), while the two denser layouts (60/90 and 40/60) were fitted with a linear regression model. The nonlinear regression model revealed that the second order polynomials were significantly non-zero for all the tail lengths (**Tab. 6**). This result suggested that for the sparser layouts increasing the letter height up to the maximum allowed by the visual angle is a good strategy to improve the reading performance. For the denser layouts, the linear regression model revealed that the slopes were not significantly different than zero (**Tab. 6**). Only for tail length 3° and layout 60/90 the slope is significantly not-zero. This result suggested that when the pixel number and density increased (i.e. the visual resolution increases) small characters can be easily recognised, and the increase in the letter height became less important.

**Table 6.** Result of the linear (y = B_0_ + B_1_x) and nonlinear (y = B_0_ + B_1_x + B_2_x^2^) regression models.

|  |  |  |  |  |  |  |  |  |
| --- | --- | --- | --- | --- | --- | --- | --- | --- |
| Tail length: 1° |  |  |  |  |  |  |  |  |
|  | 100/150 |  |  | 80/120 |  |  | 60/90 | 40/60 |
| R squared | 0.7125 |  |  | 0.4293 |  |  | 0.02253 | 0.04248 |
|  | B<sub>0</sub> | B<sub>1</sub> | B<sub>2</sub> | B<sub>0</sub> | B<sub>1</sub> | B<sub>2</sub> | Slope / B<sub>1</sub> | Slope / B<sub>1</sub> |
| Best-fit values | -150.9 | 77.51 | -6.299 | -77.02 | 59.43 | -5.117 | 1.946 | -2.865 |
| p value | < 0.0001 | < 0.0001 | < 0.0001 | 0.0311 | 0.0002 | 0.0311 | 0.2982 | 0.1510 |
| F | 24.61 | 36.28 | 24.61 | 4.938 | 16.44 | 4.938 | 1.106 | 2.130 |
| DFn | 1 | 1 | 1 | 1 | 1 | 1 | 1 | 1 |
| DFd | 47 | 47 | 47 | 47 | 47 | 47 | 48 | 48 |
| Tail length: 2° |  |  |  |  |  |  |  |  |
|  | 100/150 |  |  | 80/120 |  |  | 60/90 | 40/60 |
| R squared | 0.7261 |  |  | 0.5020 |  |  | 0.03924 | 0.04495 |
|  | B<sub>0</sub> | B<sub>1</sub> | B<sub>2</sub> | B<sub>0</sub> | B<sub>1</sub> | B<sub>2</sub> | Slope / B<sub>1</sub> | Slope / B<sub>1</sub> |
| Best-fit values | -151.4 | 71.63 | -5.824 | -109.2 | 64.90 | -5.498 | 2.503 | -2.853 |
| p value | < 0.0001 | < 0.0001 | < 0.0001 | 0.0029 | < 0.0001 | 0.0005 | 0.1679 | 0.1394 |
| F | 31.10 | 38.90 | 26.07 | 9.898 | 19.56 | 14.23 | 1.960 | 2.259 |
| DFn | 1 | 1 | 1 | 1 | 1 | 1 | 1 | 1 |
| DFd | 47 | 47 | 47 | 47 | 47 | 47 | 48 | 48 |
| Tail length: 3° |  |  |  |  |  |  |  |  |
|  | 100/150 |  |  | 80/120 |  |  | 60/90 | 40/60 |
| R squared | 0.7170 |  |  | 0.5252 |  |  | 0.1290 | 0.02389 |
|  | B<sub>0</sub> | B<sub>1</sub> | B<sub>2</sub> | B<sub>0</sub> | B<sub>1</sub> | B<sub>2</sub> | Slope / B<sub>1</sub> | Slope / B<sub>1</sub> |
| Best-fit values | -123.8 | 56.50 | -4.432 | -86.88 | 49.72 | -3.985 | 4.214 | -2.294 |
| p value | < 0.0001 | < 0.0001 | < 0.0001 | 0.0065 | 0.0004 | 0.0032 | 0.0104 | 0.2839 |
| F | 25.62 | 29.82 | 18.60 | 8.097 | 14.82 | 9.653 | 7.111 | 1.175 |
| DFn | 1 | 1 | 1 | 1 | 1 | 1 | 1 | 1 |
| DFd | 47 | 47 | 47 | 47 | 47 | 47 | 48 | 48 |

**Figure 6.**
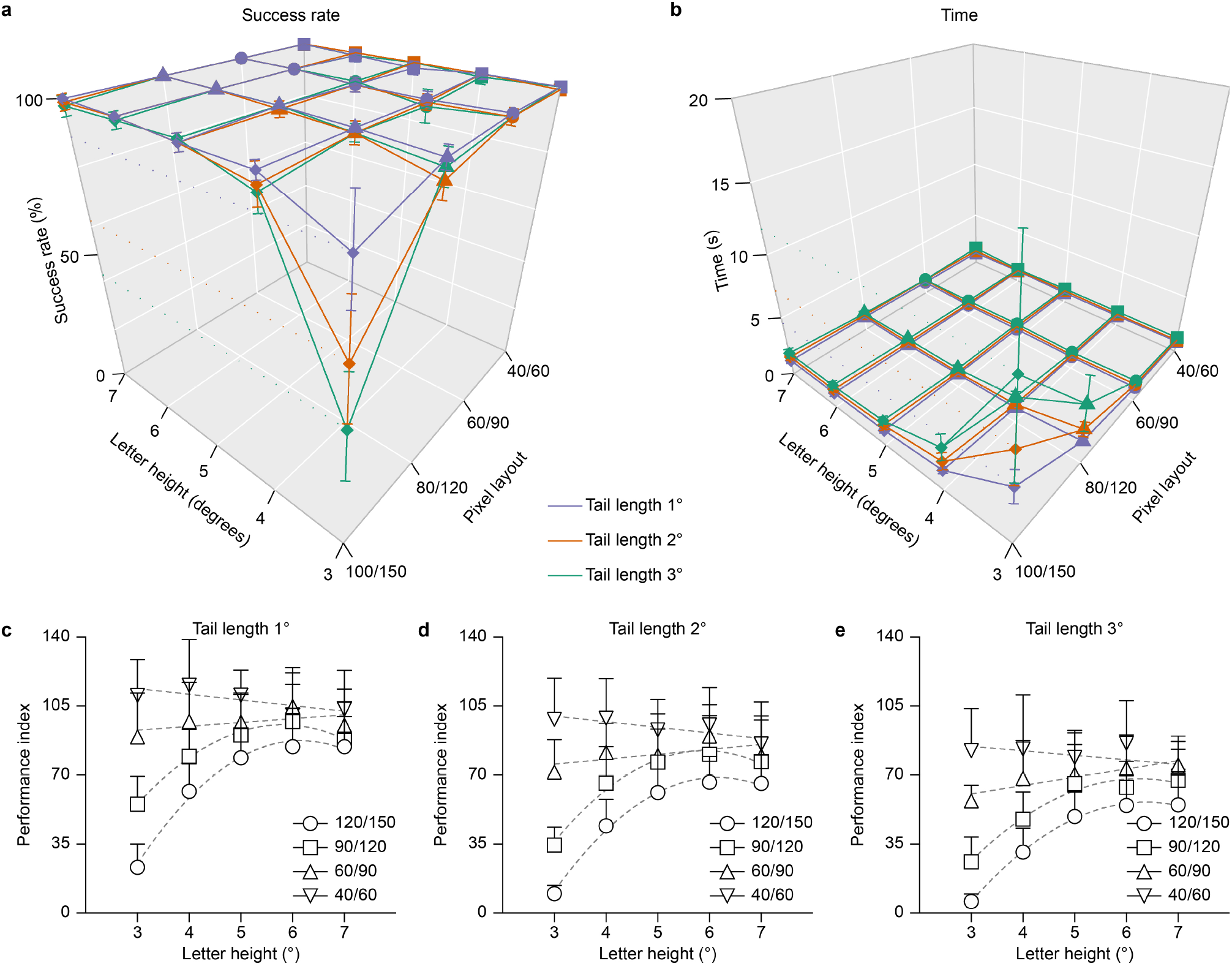
(a) Success rate (mean ± s.d., *n* = 10 participants) as a function of the letter height, pixel layout and tail’s length. (b) Time to provide a correct answer (mean ± s.d., *n* = 10 participants) as a function of the letter height, pixel layout and tail length. (c-e) Performance index (mean ± s.d., *n* = 4 participants) as a function of the letter height for the four pixel layouts. The grey dashed lines show the linear and nonlinear (second order polynomial) regression models.

The ‘word reading’ experiment demonstrated that, similar to object recognition, the visual angle has the strongest effect on performance, in particular when the pixel layout does not allow for extremely high resolution. A large visual angle allows words to be presented with a larger letter’s height, which is a good strategy to improve reading performance.

### Step perception

The third test assesses the capability to identify shallow steps, which would be useful for safe ambulation. In this cohort of participants (*n* = 11, participants 5 to 11, 13 and 15 to 17), images were converted using the 100/150 layout and four tail lengths: 0, 1, 2, 3 °. The tail length equal to zero correspond to the SB model. The participant’s visual angle was restricted to various visual angles: 5, 10, 15, 25, 35, and 45 °. Four differences in step height were tested: 10, 20, 30 and 40 cm. For each configuration (tail length, visual angle and step height) 4 repetitions were randomly presented for a total of 384 different conditions per participant (**Fig. 7a**). The participants were encouraged to take a break during the experiment to reduce fatigue. The participants were instructed to answer only if they were convinced, they knew the correct answer, otherwise to say ‘I do not know’ and the trial considered as failed. Similar to previous tests, both the percentage of correct answers (**Fig. 7b**) and the time required to provide a correct answer (**Fig. 7c**) showed that the increase of the visual angle resulted in a higher success rate and lower time to response. The performance index (**Fig. 7d**) were computed with a chance level of 50%, therefore the performance index was set to 0 when the success rate was 0%, as well as when the performance index was negative (i.e. success rate below the chance level). For each tail length, a nonlinear regression model (second order polynomial) revealed that the second order polynomials are significantly non-zero (slope, B_1_ coefficient) thus suggesting that the visual angle significantly affected the performance index (**Tab. 7**). Polling the data together, one curve cannot fit all the data sets (p = 0.0134, F = 2.380, DFn = 9, DFd = 252) suggesting that the tail length also affected the performance index.

**Table 7.**
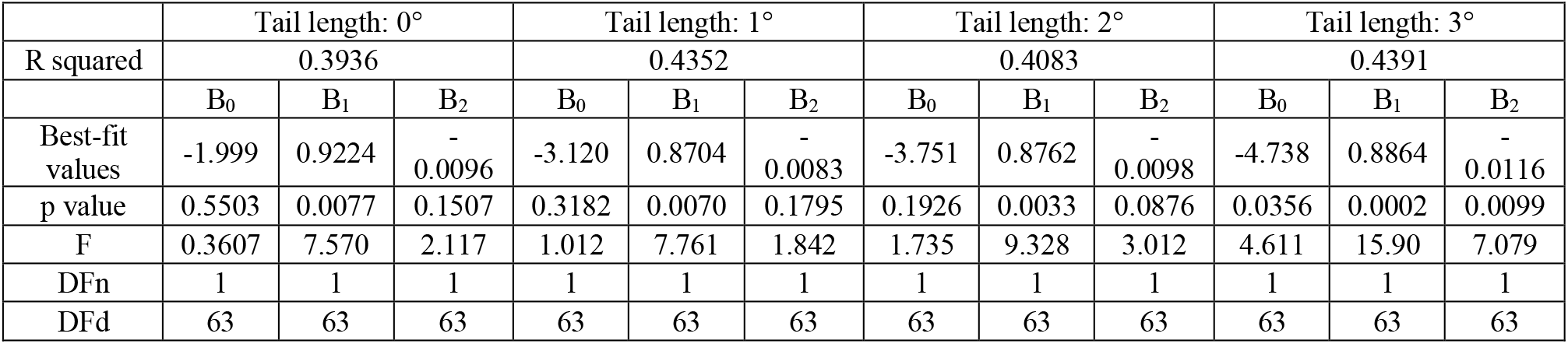
Result of the nonlinear regression model: y = B_0_ + B_1_x + B_2_x^2^.

**Figure 7.**
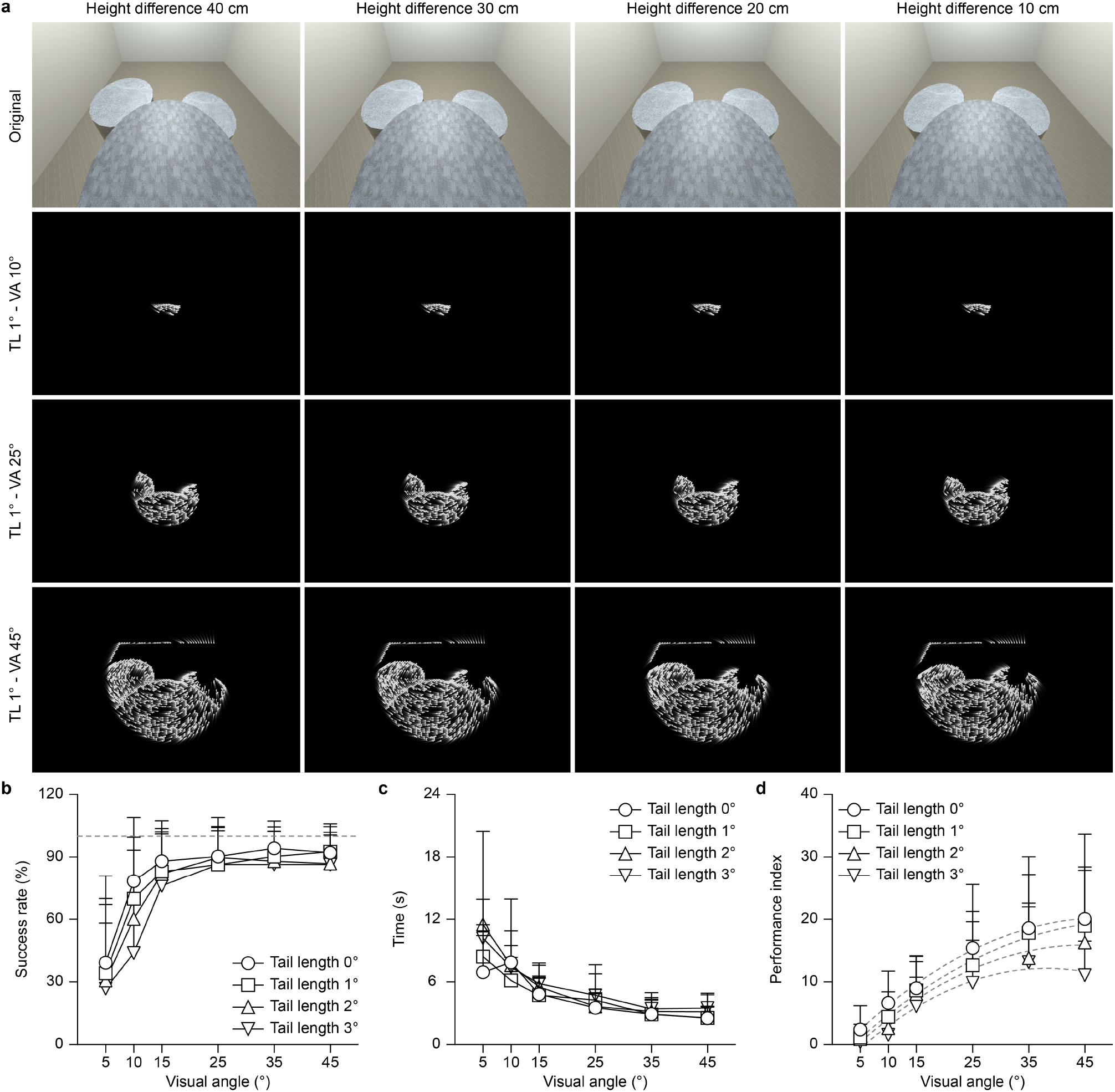
(a) Rendering of the ‘step perception’ test as a function of the difference in height (columns) and the visual angle (VA, rows). The first row corresponds to the original view. Only 3 visual angles (10 °, 25 ° and 45 °) are shown for a tail length (TL) of 1 °. (b) Success rate (mean ± s.d., *n* = 11 participants) as a function of the visual angle for the four tail lengths. The grey dashed line shows a success rate of 100%. (c) Time to provide a correct answer (mean ± s.d., *n* = 11 participants) as a function of the visual angle for the four tail lengths. (d) Performance index (mean ± s.d., *n* = 11 participants) as a function of the visual angle for the four tail lengths. The grey dashed lines show the nonlinear regression model (second order polynomial).

The ‘step perception’ experiment confirmed that the visual angle has a strong effect for the increase of the performance.

### Street crossing

The last test is a representation of typical outdoor daily activity which requires orientation capability in a dynamic environment. In this cohort of participants (*n* = 9, participants 7 to 11, 13 and 15 to 17), images were converted using the 100/150 layout and four tail lengths: 0, 1, 2, 3 °. The tail length equal to zero corresponds to the SB model. The participant’s visual angle was restricted to various visual angles: 10, 15, 25, 35, and 45 °. The smallest visual angle (5 °) was removed because it was not possible for participants to perform the test. For each configuration (tail length and visual angle) 6 repetitions were randomly presented for a total of 120 different conditions per participant (**Fig. 8a-c**). The participants were instructed to safely cross the street regardless of the time taken. Since the gaps were randomly presented, we quantified only the percentage of correct passages (**Fig. 8b**) and not the time required. For each tail length, the linear regression model revealed that the slopes are significantly non-zero for tail lengths 0 °, 2 ° and 3 °. The slope is not significantly different than zero for the tail length 1 °, for which the linear regression does not fit well the data (**Tab. 9**).

**Table 9.** Result of the linear regression model: y = B_0_ + B_1_x.

| Table 9. Result of the linear regression model: $y = B_0 + B_1x$. | | | | |
| --- | --- | --- | --- | --- |
|  | Tail length: 0° | Tail length: 1° | Tail length: 2° | Tail length: 3° |
| Slope / $B_1$ | 0.6640 | 0.3907 | 0.7136 | 1.061 |
| R squared | 0.1796 | 0.05728 | 0.1580 | 0.3848 |
| p value | 0.0037 | 0.1133 | 0.0069 | < 0.0001 |
| F | 9.416 | 2.613 | 8.068 | 26.90 |
| DFn | 1 | 1 | 1 | 1 |
| DFd | 43 | 43 | 43 | 43 |

**Figure 8.**
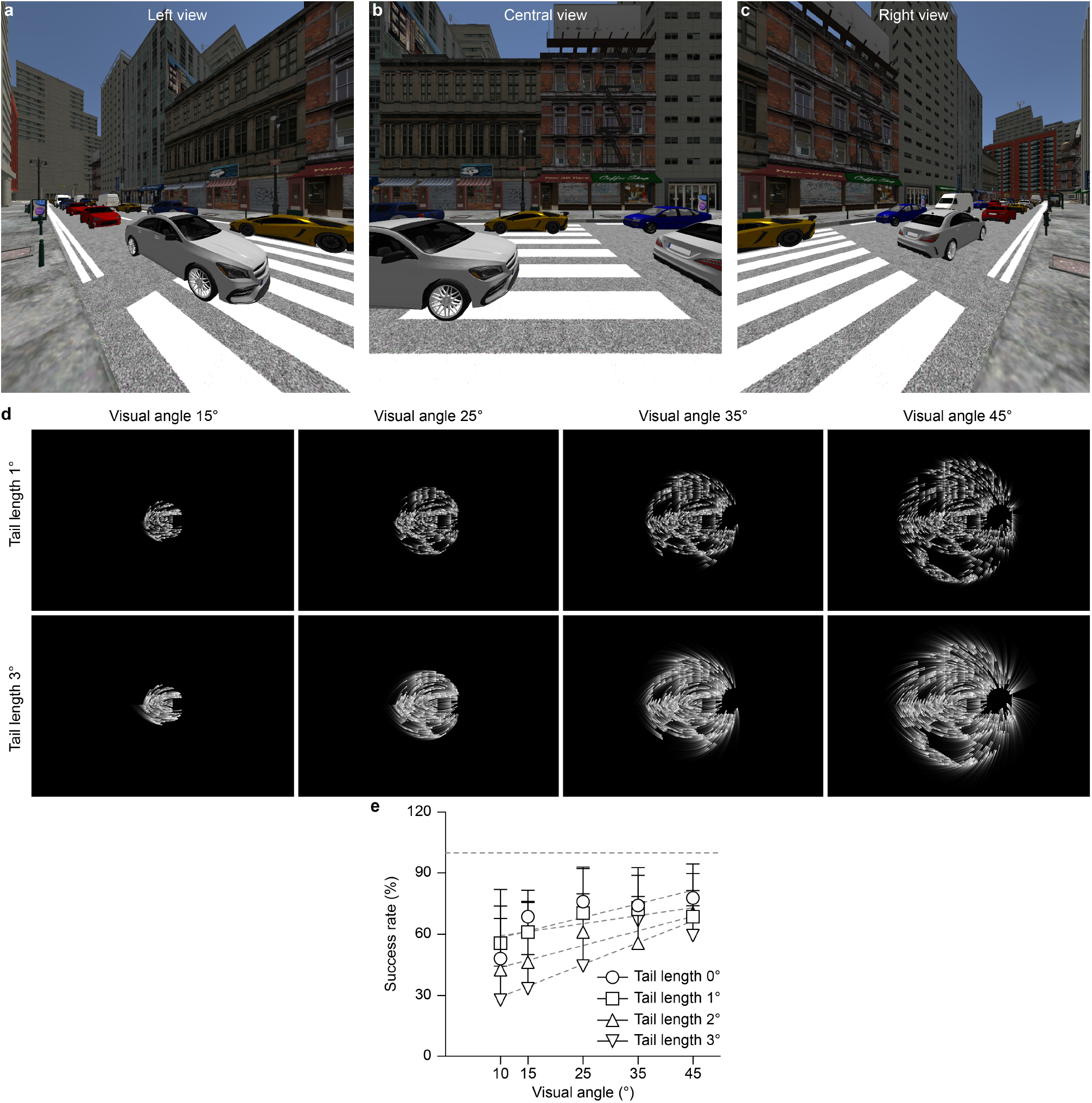
(a-c) Original view of the ‘street crossing’ test with three views: left view (a), central view (b) and right view (c). (d) Rendering of the ‘street crossing’ test (left view) as a function of the visual angle (columns) and the tail length (rows). Only four visual angles (15 °, 25 °, 35 ° and 45 °) and two tail length (1 ° and 3 °) are shown. (e) Success rate (mean ± s.d., *n* = 9 participants) as a function of the visual angle for the four tail lengths. The grey dashed line shows a success rate of 100%. The grey dashed lines show the linear regression model.

Polling the data together, the differences between the slopes are not statistically significant (p = 0.7074, F = 1. 443, DFn = 3, DFd = 172). Since the slopes are not significantly different, it is possible to calculate one slope for all the data (B_1_ = 1.430). However, the line elevations (B_0_) showed a statistically significant difference (p < 0.0001, F = 12.15, DFn = 3, DFd = 175) suggesting that the tail length affected the success rate.

As for the previous tests, also the ‘street crossing’ test highlighted the importance of the visual angle for users of retinal prostheses.

## Discussion

In the past decade, several studies assessed the performance of participants under pixelated vision using either HMDs and computer monitors.

A first subset of studies focused mostly on resolution and the capability to recognise faces, objects or text[27,29–32]. In these studies, the pixel number (m x n) was the parameter addressed the most, within a range from 8 × 8 to 40 × 50, and the effect of grey levels and pixel dropout was also evaluated. A common result among the studies is the increase in performance with an increase in the number of pixels. Remarkably, the visual angle was consistently not taken into account, even if it was not a fixed parameter since it was changing with the pixel number: if a 20 × 20 array would provide a 7 × 7 degrees of visual angle, a 34 × 34 array would provide a 12 × 12 degrees of visual angle[32]. In this latter report, the reading speed was found to decrease with the reduction of the visual angle. Other studies focused more on navigation skills and obstacle avoidance[8,9,33]. Here, the visual angle become an important factor to improve behavioural performance. While both approaches (maximisation of the pixel number or the visual angle) are currently adopted respectively to improve of recognition skills or mobility skills, the development of large-field and high-density devices like POLYRETINA opens up the possibility to combine both features in a single device.

With conventional wired retinal prostheses, the limiting parameter was always the maximum number of electrodes which could fit in the device due to the presence of the implantable pulse generator, the efficiency of the wireless power transfer and the size of the trans-scleral cable. Therefore, while keeping the electrode number constant, it might be preferable to increase the electrode density at expenses of the visual angle since a large retinal coverage with poor electrode resolution will result in an increasing difficulty to recognise objects and shapes, which is essential also for safe navigation. The advantage of a photovoltaic prosthesis like POLYRETINA is the capability to include a theoretically unlimited number of pixels, and therefore to scale the number of pixels proportionally to the visual angle. Under such conditions, it appeared that the visual angle played a major role for object recognition, safe navigation and other common daily activities. Indeed, our results showed that the visual angle is consistently the most significant factor to increase behavioural performance.

Another important parameter is the tail length. It is intuitive that the distortion of the phosphene’s shape impairs the behavioural performance. This distortion is caused by the activation of the axon of passages, which is a main limitation of epi-retinal prostheses. However, several approaches were recently reported to overcome the direct activation of retinal ganglion cells and favour the network-medicated activation, mainly based on waveform shaping[34,35] and input filters[36]. In humans, it was also demonstrated that long sinusoidal-like pulses promote the perception of circular phosphenes[37]. Improving the epi-retinal stimulation to avoid distorted phosphenes is a fundamental step to improve performance during daily activities.

## Conclusion

We believe that profoundly and totally blind patients will benefit from the restoration of artificial vision within a substantially large visual angle. However, the visual angle cannot be decoupled from the pixel count and density. Wide-field and high-density photovoltaic prostheses possess the advantage of restoring a large visual angle with a moderate pixel resolution, and thus become a helpful and valuable visual aid to patients with retinitis pigmentosa. Experiments with simulated prosthetic vision might help in designing the layout of wide-field and high-density to be tailored to the patient’s needs. However, results from these models must be considered carefully, since the model cannot recapitulate the complex psychophysical situation that implanted patients experience. For instance, the model implemented in this work considered the irregular shape of the phosphenes, which is relevant for those devices aiming at direct stimulation of retinal ganglion cells. However, other known phenomena, such as the desensitization of the retinal ganglion cells activity upon repetitive stimulation during network-mediated stimulation, are not yet implemented. Desensitization is an important parameter to be considered in those devices, like POLYRETINA, which relies mostly on the network-mediated stimulation.

## Acknowledgement

We would like to acknowledge the The Virtual Reality Facility at Campus Biotech in Geneva. This work was supported by École polytechnique fédérale de Lausanne, Medtronic and The Fondation E. Et G. Gelbert.

## Author contributions

J.T.T. wrote the code and ran the tests. E.M. participated in writing the code. D.G. designed and led the study, and wrote the manuscript. All the authors read and accepted the manuscript.

